# Distributed neural computation and the evolution of the first brains

**DOI:** 10.1101/2025.10.03.680388

**Authors:** Vikram Chandra, Mehrana R. Nejad, Allison P. Kann, Ananya Salem, Karl A.P. Hill, Richard Schalek, Jeff Lichtman, L. Mahadevan, Mansi Srivastava

## Abstract

The origin of brains in the Precambrian was a landmark in animal evolution, enabling new behavior and life histories. Brains likely evolved from diffuse nerve nets, but we do not know what the first brains looked like or how they were organized. Acoel worms, the likely sister lineage to all other animals with brains, offer a unique window into this transition. Here, we studied the acoel worm *Hofstenia miamia*, a marine predator that hunts planktonic invertebrates and displays other sophisticated behavior. We found that *H. miamia* has an unusual ‘diffuse brain’: a subepidermal network of dense neuropil exhibiting little regionalization or stereotypy in gross anatomy or distribution of neural cell types. Remarkably, we found that behavior in *H. miamia* is robust to large, arbitrary amputations of brain regions, suggesting that most regions can perform most computations. More brain tissue improves performance, especially on challenging tasks, but no specific brain region is required. These results lead us to propose that *H. miamia*’s brain is composed of computationally pluripotent “tiles” that interact to generate coherent behavior. This architecture suggests a trajectory for nervous system evolution in which early brains may have arisen through the condensation of diffuse nerve nets into unregionalized brains, with regionalization evolving secondarily.

## Main text

The evolution of brains was a milestone in the history of animals: it enabled new behavior, including early predatory lifestyles, and is thought to have triggered an evolutionary arms race between hunters and hunted that shaped the Cambrian explosion and modern animal diversity^1–5^. Modern animal brains are the result of this ‘Cambrian information revolution’^5,6^. Although they vary substantially in their organization and complexity, brains are centralized computing organs that serve as integrating centers; they collect sensory inputs and generate coherent organismal behavior^7–11^. Brains are highly regionalized, with specific regions specialized for specific tasks^12,13^. They are also stereotyped; within a species, all animals have the same brain organization. Brains evolved in the Bilateria (bilaterally-symmetric animals) some 550-600 million years ago, likely from ancient diffuse nerve nets^9,14,15^. Indeed, the first animal nervous systems were likely diffuse nets, and extant cnidarians (and possibly ctenophores) retain a version of diffuse nets^9,16^. Such nervous systems do not possess the core features of brains; for instance, they are not regionalized or stereotyped^17–19^. Instead, these features likely evolved during or after the origins of centralized nervous systems in bilaterians.

Over the last two centuries, much has been learned about the organization and function of diffuse nets. In the 1870s, Romanes and contemporaries, through a series of experiments involving incisions and amputations of jellyfish, first showed that cnidarians possess an unpolarized conducting substance distributed through their bodies^17,18,20^. A century later, electrophysiology and advances in histology reinforced these findings, and more recent work has begun to reveal detailed circuit explanations for how diffuse nets compute^21–27^. However, the large number of differences between diffuse nets and centralized brains has made it difficult for biologists to imagine what the first brains may have looked like, or how nets transitioned to brains^9,11,28^.

Acoel worms are ideally poised to shed light on these questions. Acoels belong to the likely sister lineage to all Bilateria^29–32^ (Fig. 1a). Moreover, acoels are thought to possess some of the simplest brains. In other words, their nervous systems appear intermediate between diffuse nets and centralized brains^11,33–37^. Comparing acoel brains to those of other bilaterians could reveal the architecture of ancient brains. However, acoel brains and behavior are poorly understood. Here, we study an acoel, the three-banded panther worm *Hofstenia miamia* (Fig. 1b)*. H. miamia* lives in intertidal regions on beaches, mangroves, and salty lakes in the North Atlantic and Caribbean^9^. The worms are hermaphroditic and range in size from ∼500µm to 1.5cm, and have relatively simple anatomy with a blind gut and few discrete organs^31,38,39^.

**Figure 1:**
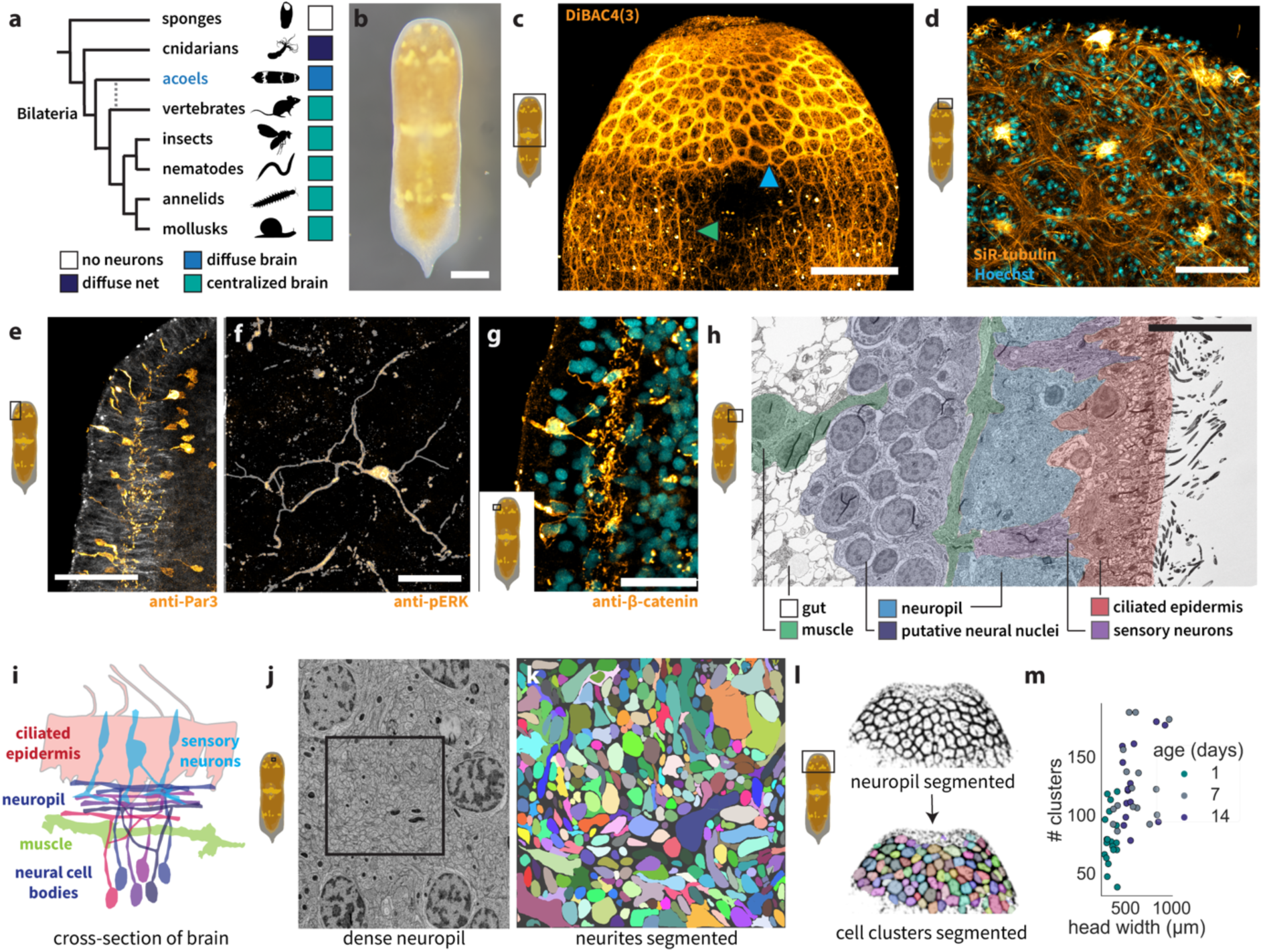
The organization of a diffuse brain. a) Simplified phylogeny of animals shows that acoel brains are likely intermediate between cnidarian diffuse nets and the centralized brains of typical bilaterians, with their alternative phylogenetic position annotated (dashed gray line). b) Photograph of juvenile *Hofstenia miamia*. c) Staining with voltage dye reveals a superficial network of dense neuropil (blue arrow) that extends into a sparser posterior nerve net (green arrow). d) Close-up view of neuropil stained sparsely with tubulin dye (orange) reveals that the neuropil (orange) contains many neurites running in parallel, with cellular clusters (cyan) interspersed between neurite bundles. Sensory neurons (likely clusters of H1 cells; bright orange) are set within many of these patches. e) Cross-section of brain stained with a Par3 antibody reveals that the brain has two layers: superficial neuropil, and deeper cell bodies that project outward. f) Staining with an ERK antibody (z-projected segmentation overlaid) shows that brain interneurons can be multipolar, with a central cell body generating multiple neurites. g) Cross-section of brain stained with an antibody against β-catenin reveals another sensory neuron class (possibly H2^43^) with two projections that innervate brain neuropil. h) Electron microscopy cross-section shows the fine organization of the brain, confirming the relative configuration of tissue types within the head. The superficial neuropil is visible immediately beneath the skin, while neural cell bodies lie deeper in the tissue, internal to body wall muscle (green). i) Schematic of cross-section of brain. j) Electron microscopy close-up of the brain shows dense neuropil; the box is a 6.7x6.7µm square. k) Segmenting neural projections within the highlighted box in (j) reveals over 400 neurites in a single section of neuropil. l) Segmentation of cellular clusters within neuropil allows quantification of brain structure and its variability. m) Quantifying the numbers of cellular clusters across brains reveals that, although cluster numbers increase with age (i.e. days after hatching) and size (i.e. head width, a good proxy for overall body size^42^), worms vary widely in how many clusters they possess. Linear regression p<0.0001, n=49. Scale bars: 1mm (b), 200µm (c), 50µm (d,e), 20µm (f,g), 10µm (h).

Taxonomic descriptions suggest that the worms are “voracious predators”^38,40,41^, and in recent work we found that they display complex reproductive behavior^42^, suggesting they have a rich and sophisticated behavioral repertoire. We developed methods for quantitative analysis of *H. miamia*’s brain and behavior and found that *H. miamia* has a ‘diffuse brain’ that lacks the regionalization and stereotypy of well-studied bilaterian brains. We identify organizing principles for this diffuse brain, and suggest a trajectory for the evolution of the first brains.

## The *H. miamia* brain lacks anatomical regionalization and stereotypy

In her taxonomic species description of *H. miamia,* Correa suggested that its brain was a subepidermal ‘ganglionic ring’ encircling the head^39^. Building upon this and recent work^36^, we used a variety of imaging techniques (live dyes, immunofluorescence, and electron microscopy) to understand the brain at high resolution.

The brain contains a superficial network of dense neuropil, i.e., a structure composed primarily of neural projections, that lies peripheral to the body wall muscle and immediately internal to the epidermis (Fig. 1c-e; Fig. S1a-c). Each ‘edge’ of the network contains neurites running in parallel. Superficial cellular patches tile the neuropil, and these often contain cells that coalesce into a sensory structure within each patch. These patches are composed of likely ‘H1’ receptors, sensory cells previously described in multiple acoels^43^ (Fig. 1d; Fig. S1d), and also contain muscle cell bodies (Fig. S1e). The cell bodies of brain neurons lie ∼30-50µm beneath the skin, internal to the body wall muscle. These cell bodies project outward into the neuropil, ∼10-20µm below the skin (Fig. 1e-h). The two ‘layers’ of the brain are primarily composed of the same interneurons, with deep cell bodies generating neurites that project peripherally and coalesce into dense, superficial neuropil (Fig. 1h,i). The brain displays a slight dorso-ventral asymmetry: there are fewer network ‘edges’ in the neuropil near the ventral midline (Fig. S1a,b), possibly to accommodate the male copulatory apparatus^42^.

With regards to cellular anatomy, we identified both unipolar and multipolar neurons in the brain (Fig. 1e,f). While the complete 3D structures of these neurons (and whether neurites are separable into distinct axons and dendrites) remain unclear, we found that they can have projections that extend well over 100µm (Fig. 1d; Fig. S1f). Sensory cells embedded in the skin project into the neuropil (Fig. 1e,g; Fig. S1g). Long cilia are visible near the mouth, likely corresponding to distinct sensory cell type(s) (Fig. S1a-c,h). Outside the brain, we detected multiple distinct innervated organs: the frontal organ (Fig. S1i), statocyst (Fig. S1j-l), penis (Fig. S1m), and pharynx (Fig. S1n). The brain itself connects to a sparse posterior network that extends through the body (Fig. 1c). Additionally, we found dense peripheral neural processes distributed through the skin (Fig. S1o; Video S1).

*H. miamia* neuropil forms a network, with superficial resemblance to the diffuse nets of cnidarians and ctenophores. To understand the internal structure of *H. miamia* neuropil, we used electron microscopy. We found that every region of neuropil contains hundreds of fine, reticulated neural projections (Fig. 1j,k). This density is suggestive of complex connectivity and the likelihood that this brain harbors a diversity of circuit motifs. Importantly, this organization is fundamentally different from cnidarian or ctenophore nerve nets, which are typically 2-3 neurites wide^22,44,45^, rather than hundreds or thousands of neurites wide.

Together, these results describe the structure of the *H. miamia* brain. Most remarkably, we found that this brain is virtually devoid of internal anatomical regionalization: within the neuropil, we detected no distinct regions, lobes, or stereotyped commissures. Regionalization is a core feature of typical bilaterian brains: across phyla as diverse as mollusks, arthropods, nematodes, vertebrates, and annelids, different brain regions have different structures and functions, and these regions are in the same relative positions across individuals within a species (and often over hundreds of millions of years)^12,46–48^. Given its lack of regionalization, the *H. miamia* brain appears to be fundamentally different. Next, we asked whether this brain displayed any stereotypy in its structure.

To test for stereotypy, we quantified brain structure by segmenting and counting the sensory patches that tile the neuropil (Fig. 1c,d,l). We found that as worms grow during juvenile development, the number of these patches increases (Fig. 1l,m). However, even among worms of the same age, size, and rearing environment, we found an order of magnitude of variation in the number of patches within their brains (Fig. 1m). Thus, while every brain has similar network topology, every brain is also different in its precise shape. This indicates that the same neural circuits cannot be in the same relative locations across worms. In other words, the *H. miamia* brain, unlike previously-studied bilaterian brains, appears to lack regionalization and stereotypy, and it suggests that this brain must be organized differently.

## *H. miamia* behavior is robust to large, arbitrary amputations of brain tissue

The lack of anatomical regionalization within the *H. miamia* brain suggests that function may not be regionalized either. Taking advantage of *H. miamia*’s ability to survive substantial injury, we sought to assess functional brain regionalization by amputating brain regions and studying subsequent impacts on behavior.

To study the consequences of removing brain regions, we first developed an ethologically-relevant behavioral assay. *H. miamia* are thought to forage by hunting other small marine invertebrates in the wild, and they eat brine shrimp or rotifers in the lab. We developed a foraging assay in which worms are presented with a choice between a block of agarose containing prey rotifers and a control block of agarose containing no prey. We found that hungry worms navigated to the rotifers and attempted attacks (Video S2), while continuously fed (Fig. 2a-c) worms did not. Next, to ask whether this foraging behavior ceased when the worms had consumed enough food: i.e., to ask whether hunger and satiety are opposing motivational forces that regulate foraging behavior, we first starved worms and then fed them for 6h prior to studying foraging. These sated worms did not forage, showing that foraging behavior in *H. miamia* is motivation-dependent as in typical bilaterians^49^ (Fig. S2a).

**Figure 2:**
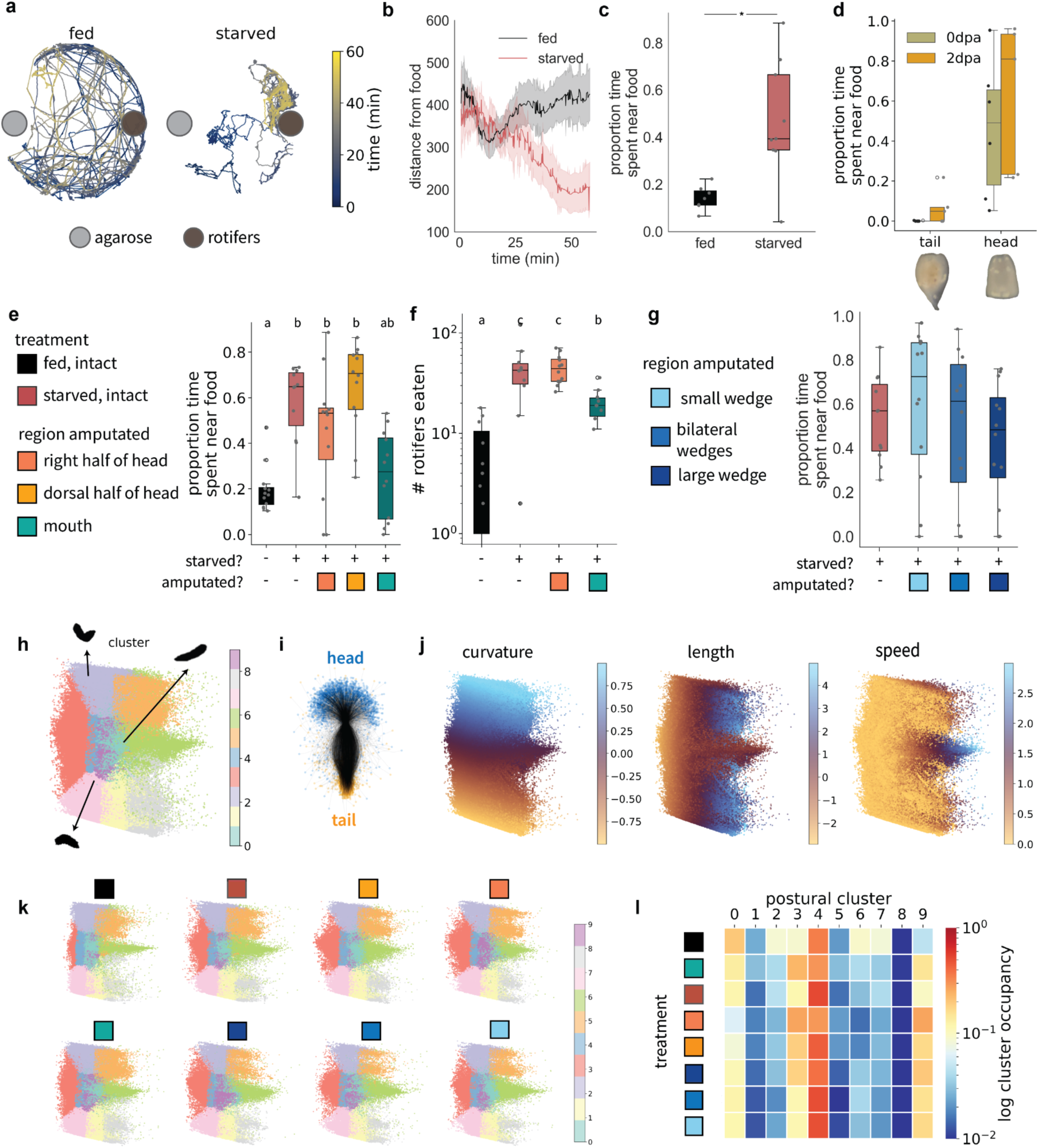
***H. miamia*’s foraging behavior is robust to brain amputations.** a) Examples of tracks for a representative fed and starved worm presented a choice between agarose blocks with and without rotifers. Trajectories colored by time. b) Starved worms approach the rotifers; fed worms generally do not (see also Fig. S3a). Error band represents standard error. c) Quantifying this behavior as the proportion of time spent near food provides a robust metric of foraging behavior (see Methods); t-test FDR-corrected p=0.03,n>=6, data also used in Fig. S3a. d) Amputated tails do not forage, either immediately (0 dpa: t-test p=0.01, n>=5) or two days after (2dpa: t-test p=0.01, n>=5) amputation, while amputated heads do. e) Amputation of the right or dorsal half of the head does not affect foraging behavior, while amputation of the mouth inhibits foraging. f) Quantifying the number of rotifers in the guts of worms shows that amputation of half the head does not affect feeding. Letters above boxplots in (e) and (f) indicate significance levels after one-way ANOVA and Tukey post-hoc tests; corrected ps<0.05. g) Removing tissue of varied size and geometry from the brain also does not affect foraging behavior. One-way ANOVA p=0.54, n>=10. h) A low-dimensional postural space (i.e. of the first two principal components of the space defined by curvature, length, and speed) captures major elements of *H. miamia*’s postures during foraging, with k-means cluster identities shown, and representative worm postures overlaid for some clusters. See Table S1 for mean values of curvature, length, and speed for each cluster. i) Overlaid and aligned splines fitted to tracked worm midlines from a representative worm show the space of possible postures - and especially curvature - for a worm. j) Normalized curvature, worm length (z-scored within worm), and speed appear as gradients within the postural space. k) Projecting amputated worm postures into this postural space shows that irrespective of amputation, all worms express all elemental postures. l) Comparing postural cluster occupancies shows that fed, intact worms are most different in their postural behavior, while amputated worms generally display only minor differences in occupancy compared to starved, intact worms. Legend for treatment colors in (e) and (g). See Tables S2 and S3 for statistics.

Next, we asked how foraging behavior was affected by amputating brain regions. First, we performed a transverse amputation to separate heads from tails in starved worms. We found that tail fragments, which lack brains, did not forage and were unable to move (Fig. S2b), while head fragments foraged normally both immediately after amputation and two days later (Fig. 2d), when their wounds had closed but before their tails had regenerated^31,50^. This shows that the posterior nerve net does not independently generate coherent movement or induce goal-directed behavior, and that the brain is likely necessary for coordinated behavior. We then removed the right half of the head (Fig. S2c). Worms missing half their heads did not display obvious movement defects (Fig. S2d). More importantly, we found that two days after amputation, worms missing half their heads foraged similarly to intact worms, when the wound had closed but new tissue had not yet emerged (Fig. 2e; Fig. S2e-g,i,j). Similarly, removing the dorsal half of the head, or the left half of the head, did not affect foraging two days after amputation (Fig. 2e; Fig. S3a).

This robustness to amputation could result either from the inherent structure of the *H. miamia* brain, or through rapid regeneration after injury. We found that worms missing half their heads foraged successfully immediately after amputation (Fig. S3b), suggesting that foraging behavior does not require regeneration of missing brain tissue. To confirm this finding, we irradiated worms, ablating their pluripotent stem cells and thus inhibiting regeneration^31^. Worms without stem cells foraged successfully regardless of whether their heads were intact or had been amputated (Fig. S3c). These data show that robustness to amputation is an intrinsic property of the *H. miamia* brain.

We next asked whether this robustness extended beyond the specific computations and movements involved in the approach phase of foraging. To study the capture phase of foraging, we provided worms access to a controlled density of free-moving rotifers for a defined time period, and quantified the number of rotifers they consumed. Worms missing half their heads consumed similar numbers of rotifers to intact starved controls (Fig. 2f). These data show that amputation of large fractions of the brain do not prevent worms from executing the set of tasks necessary for foraging. In addition, these amputations include conditions in which the statocyst, frontal organ and associated neurons were damaged or entirely removed, showing that these organs are dispensable for all aspects of foraging behavior.

Next, to further test the generality of brain robustness, we removed wedges of differing sizes from the heads of worms: a small wedge, two bilateral wedges (to account for the possibility of bilaterally-symmetric, specialized regions with redundant roles), and a large wedge that removed the majority of the head (Fig. S3d; Video S3). We found that worms foraged robustly despite the loss of these brain regions (Fig. 2g). In many of these conditions, wound closure involves the attachment of dorsal and ventral wound edges^50^, likely connecting brain regions that are normally separate (Fig. S2g). Despite these abnormal structural configurations, worms displayed robust foraging behavior. Moreover, analysis of the angular distributions of the trajectories of these worms found no major differences across treatments, and no turn bias, even after large lateralized amputations (Fig. S3e). This shows that most or all regions of the *H. miamia* brain can locally perform motor control. Finally, we also conducted a wide variety of additional amputations, removing wedges of varying size and location from worm heads. These also did not affect foraging behavior (Fig. S3f).

While removing large amounts of brain tissue did not affect foraging, we found that removing the mouth did inhibit successful foraging (Fig. 2e; Fig. S2h). These worms have no obvious movement defects (Fig. S2d). Worms missing their mouths consumed fewer rotifers than intact worms or those missing half their heads, but significantly more than intact fed worms did (Fig. 2f), suggesting that the mouth is not strictly necessary to assess internal nutritional state, or to coordinate goal-directed appetitive or consummatory foraging behavior. Together, these data suggest that amputation of the mouth may result in a chemosensory-specific defect, consistent with the presence of specific sensory cell types near the mouth (Fig. S1a-c,e)^43^, and indicate that robustness is a feature of brain computations downstream of chemosensation.

Overall, these data show that behavior in *H. miamia* is remarkably robust to the removal of large, arbitrary majorities of brain tissue. Nonetheless, it is possible that worms with amputated brain regions experience some subtle behavioral defects. We reasoned that these hypothetical defects may be observable in their motor coordination, since the worms were clearly able to detect prey, to sense their internal state, and to select actions necessary for foraging. We therefore quantified worm posture during foraging at high resolution, by tracking keypoints on the worm’s midline. Using these data, we generated a low-dimensional postural space during foraging from starved worms with intact brains (Fig. 2h-j; see Methods). The boundaries of the occupied region of this space (Fig. S4a) reveal core biomechanical constraints on *H. miamia*’s movement. *H. miamia* has two motor systems: epidermal cilia (Fig. 1h, Fig. S1h) polarized along the anterior-posterior axis beat to produce thrust and allow forward gliding, and distributed groups of muscle (including an orthogonal grid of body wall muscle) contort to allow turning (Fig. 1h)^51^. Worms in sharp turns experience no forward displacement, likely because the cilia in different body regions are beating in different directions. Worms are fastest when at intermediate length, but stretch and compress themselves frequently (Fig. S4a).

Next, we used k-means clustering to segment elemental postural states within this space (Fig. 2h; Table S1). We then projected onto this space similar data from fed worms with intact brains as well as starved worms subjected to numerous amputations (i.e., data from Fig. 2f,g).

We found that fed, intact worms were most different in their postural expression, consistent with their lack of foraging behavior (Fig. 2k,l; Fig. S4b,c; Table S2,3). Arbitrary amputations of brain regions did not affect the generation of postures or the transitions between postures, although we detected small but significant effects on relative occupancy of specific postural states and transitions (Fig. S4b,c; Table S2,3). In addition, our postural analyses did not detect differences in turning bias after amputations of the right half of the brain (Fig. 2k,l; Table S1,2; Video S3).

This reinforces the idea that brain regions locally generate postural states necessary for both left and right turns (along with all other motor commands), unlike in typical bilaterians where a specialized brain region commands distant motor systems, frequently in a lateralized manner^52,53^. In other words, these data suggest that most or all motor commands can be generated in any or all brain regions, control motor behavior locally, and can be transmitted body-wide.

Together, these data show that the *H. miamia* brain exhibits little functional regionalization. Instead, most or all brain regions are computationally pluripotent, i.e., most or all neural computations necessary for foraging occur in many or all regions of the brain, likely simultaneously. These computations include sensory processing and integration, internal state assessment, selecting actions, and motor control of most or all muscular and ciliary motor systems, i.e., the generation of all static postures and their sequences.

### Neural cell types lack regionalization within the brain, consistent with a tiled organization

How might such a brain be organized? We hypothesized that the brain is composed of tiles of similar neural circuits, each of which is computationally ‘pluripotent’. If this were true, we would expect these tiles to be composed of ensembles of similar neural cell types. Conversely, detecting substantial regionalization in the locations of individual cell types would disprove this hypothesis. Our previous work in *H. miamia* found that neural markers are distributed across the entire brain^54,55^. However, many of these markers are expressed in large subsets of neural cell types, and we reasoned that, as in typical brains^56,57^, markers of sparse cell types could exhibit greater regionalization. To test the tiling hypothesis more directly, we therefore sought markers of sparse cell types. We first integrated all relevant *H. miamia* single-cell RNA sequencing datasets^54,58^ (Fig. S5a), selected all neurons (Fig. S5b,c), and reclustered them to identify individual neural cell types (Fig. S5d,e). Analyzing this dataset of 15,262 cells, we identified 25 putative neural clusters with specific markers for most clusters (Fig. 3a). We selected markers likely to have functional relevance from this list. To overcome the possibility that some neural markers may not be represented in this dataset due to technical limitations, we also included evolutionarily conserved genes with likely roles in neurotransmitter synthesis or processing pathways that were previously identified^59–61^.

**Figure 3:**
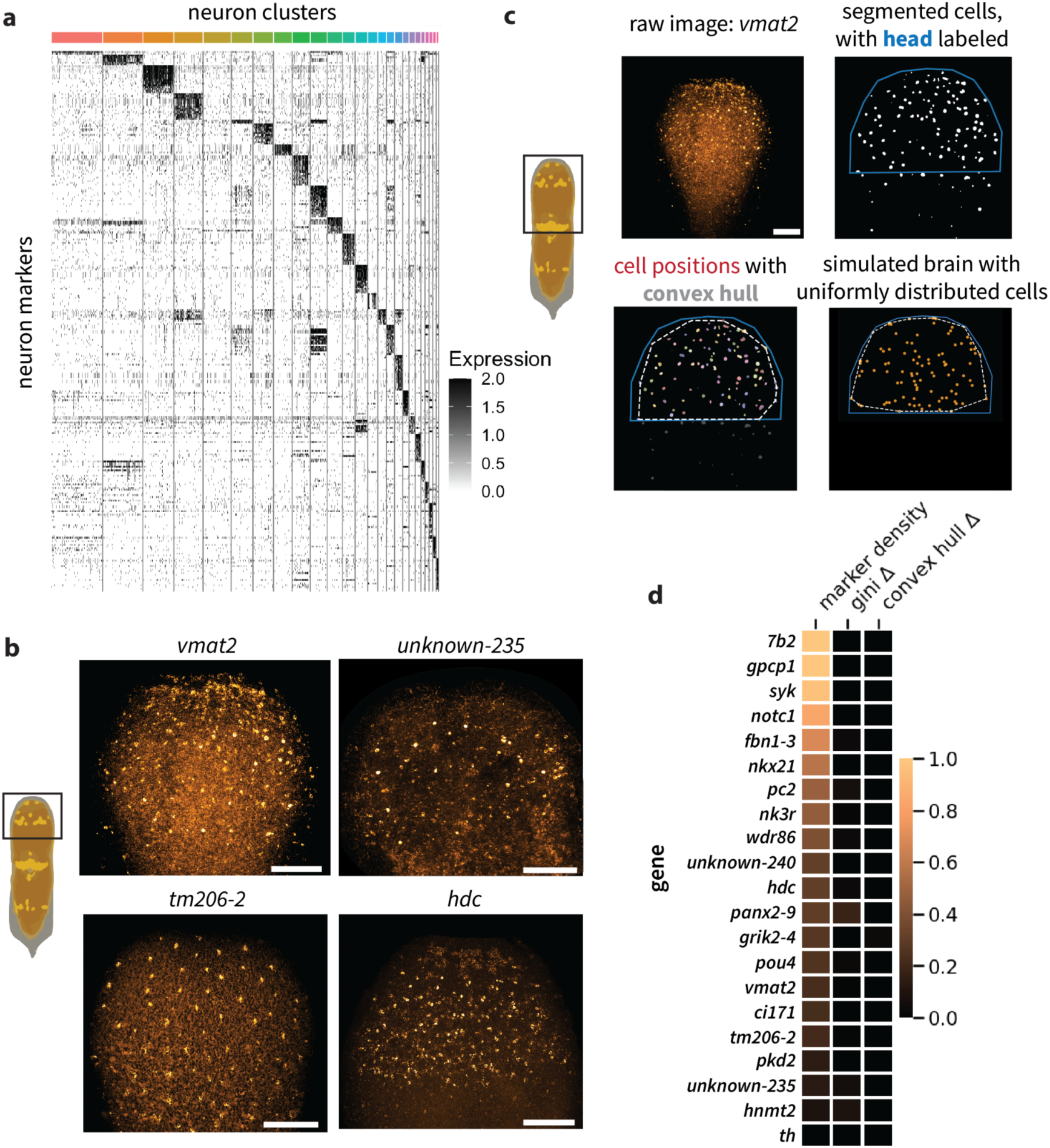
Neural cell types are distributed across the *H. miamia* brain. a) Heatmap of top 20 markers from each neural cluster shows that many markers are expressed in specific clusters, suggesting that these are cell-type specific markers. b) Fluorescence in situ hybridization of neural markers reveals that several (e.g. *vmat2*, *unknown-235*, *hdc*) label sparse, distributed populations, while others are expressed broadly (e.g. 7b2). c) Outline of analysis of spatial distributions of markers: pixel classification to segment fluorescent puncta (corresponding to cell bodies for sparse markers), fitting convex hull to segmented cells and calculating the ratio of its area to the area of the polygon describing the brain, and simulating brains with uniformly distributed cells with the same density as the empirical data. d) Neural markers exhibit little regionalization across a wide range of cell densities, as assessed by two metrics: the difference in gini coefficients for brain regions containing expression between real brains and simulated uniform brains, and the difference in the proportion of brain area containing cells between real brains and simulated uniform brains. Scale bars: 100µm.

We used fluorescence *in situ* hybridization to assess the distribution of neurons expressing these genes in worms. Of 44 genes that we screened (Table S4), we identified 21 with expression in the brain, including several that labeled sparsely-distributed neurons within the head and others that labeled large subsets of neurons (Fig. 3b; Fig. S6). Qualitatively, all identified cell populations appeared largely or entirely distributed across the brain, regardless of their density (Fig. S6). To systematically test regionalization, we quantified gene expression patterns for all markers in the head (including for previously published markers for all qualitative neural expression classes, but excluding markers of sensory neuron populations in the mouth and elsewhere in the body), and we computed two density-corrected indices of regionalization for each gene (Fig. 3c; see Methods). Irrespective of their sparseness, markers had minimal regionalization on both indices (Fig. 3d). These data are consistent with the tiling hypothesis.

Taken together, our anatomical, behavioral and molecular data all find little evidence that the *H. miamia* brain contains specialized regions. Instead, the brain may be composed of repeating, computationally pluripotent units - which we term ‘circuit tiles’.

The exact wiring architecture of circuit tiles remains unknown, but our data suggest some constraints on their organization. It is possible that the patches of sensory cells that tile the brain, with surrounding neuropil, represent computationally pluripotent circuit tiles themselves. However, some cell type markers are sparser than these (e.g. *th, hnmt2, unknown-235* - see Fig. 3d; Fig. S6). Since tiles are pluripotent, and if we assume these sparse cell types are necessary for some computation, it would follow that each tile must contain at least one representative of each cell type. Therefore the patches of sensory clusters are unlikely to be the pluripotent circuit tiles. It would further follow that the size of each tile is set by the distance between neighboring cells of the sparsest type. Alongside these constraints, it is possible that tiles vary in their exact composition and architecture. Tiles need not occupy entirely discrete regions of space; it is possible that they overlap partially. We also do not wish to imply that tiles necessarily have higher within-unit than between-unit connectivity; in fact, our anatomical studies of the brain (e.g. Fig. 1d) suggest that there is likely extensive lateral connectivity across all regions of the neuropil. Regardless, the worm’s anatomy ensures that every tile contains connections to the motor systems: epidermal cilia, peripheral, body wall, parenchymal and pharyngeal muscle. In addition, tiles likely contain one and possibly multiple sensory cells or organs, which allows each tile to locally receive some sensory input and to transmit most or all motor commands (Fig. 1i).

## On how a diffuse brain computes

The unusual organization of the *H. miamia* brain raises several questions. For instance, why does it contain multiple circuit tiles, when fewer, or perhaps even one, should do? We reasoned that while no specific brain region may be necessary for any computation, more circuit tiles may allow better performance. To seek associations between available brain area and behavior, we first reanalyzed the experiment in which we amputated different amounts of brain tissue (Fig. 2g). Analyzing these data as a function of approximately how much brain tissue was removed (rather than as a set of discrete perturbations), we found that although no amputation treatment prevented worms from foraging successfully, worms missing more brain tissue took slightly longer to find immobilized rotifers (Fig. 4a; Fig. S7a). This was not because the worms were mechanically impaired or took longer to travel the same distance (Fig. S7b); instead, worms missing more brain tissue may be worse at locating prey. To test whether this effect was reproducible, we diluted the prey stimulus by reducing the concentration of immobilized rotifers presented to the worms, making it more challenging for worms to locate prey. Worms missing ∼80% of their brains successfully located rotifers when presented a high concentration, but not when presented a low concentration (Fig. 4b), suggesting that more brain tiles may allow greater perceptual resolution and therefore behavior resulting in higher fitness. These results also showed that presenting the worms with challenging tasks can reveal subtle effects of amputation.

**Figure 4:**
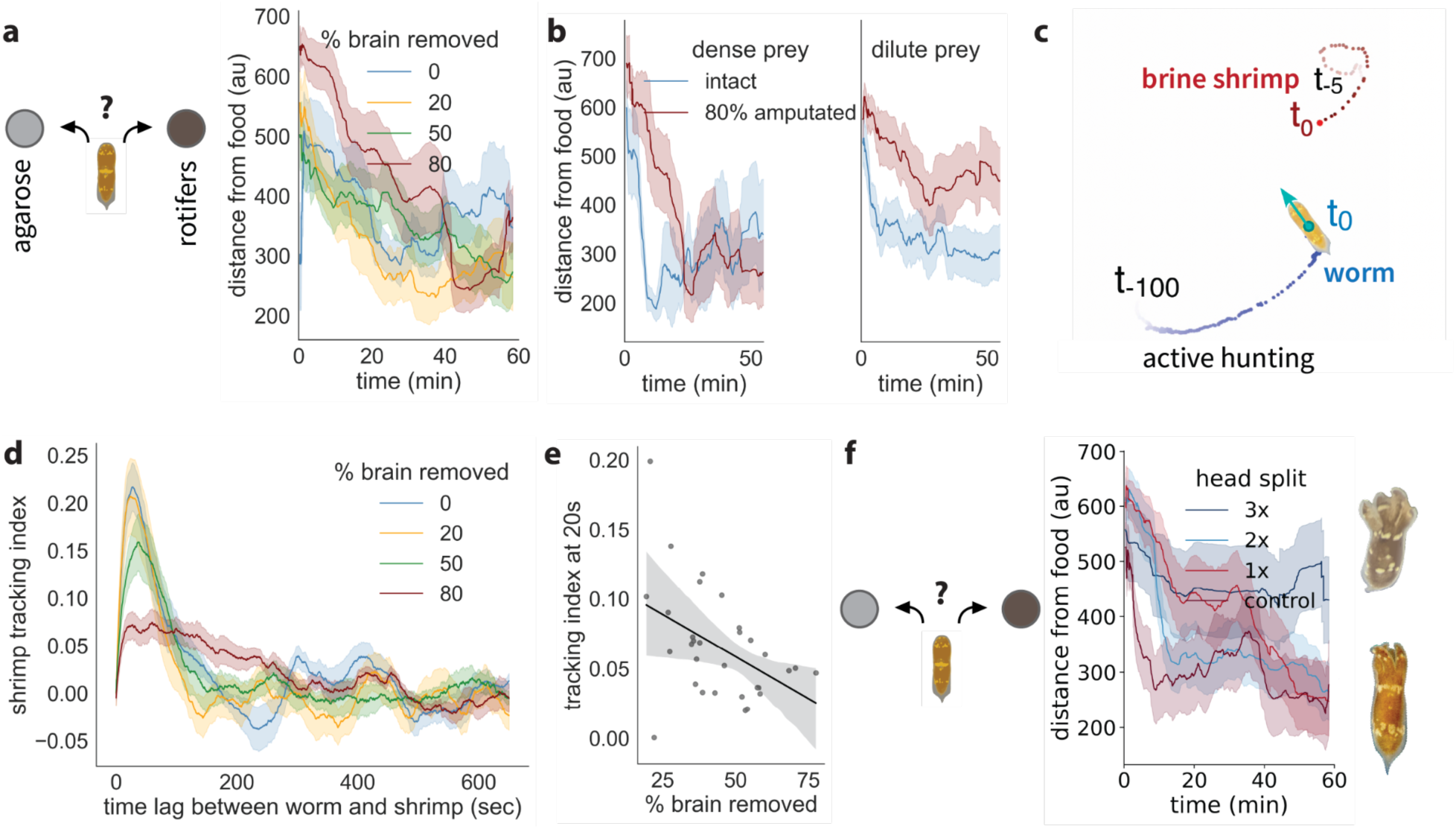
The collective behavior of circuit tiles. a) Worms missing greater fractions of their brain take longer to locate rotifers embedded in agarose (schematic shows assay design: worms choose between agarose containing rotifers and plain agarose, as in Fig. 2). b) Reproducing this effect, worms with ∼80% of their brain missing are unable to detect a low density of rotifers embedded in agarose, and are slower to detect rotifers at high density. High-density prey, intact vs amputated t-test for proportion of time spent near food: p=0.8, n=5 per treatment. Low-density prey, intact vs amputated t-test for proportion of time spent near food: p=0.02,n>=14 per treatment. c) Schematic of hunting behavior assay, showing joint tracking of worm and its prey shrimp. The recent trajectory of the shrimp is shown in red (5s); the recent trajectory of the worm is shown in blue (100s). The cyan arrow shows the direction of motion of the worm. d) Worms track shrimp positions; worms missing more brain tissue display poorer tracking. Linear regression of max. tracking index: p<0.0001, n=38. e) Worms missing more brain tissue display poorer tracking, even when their mouths are completely intact. Linear regression of tracking index at a lag of 20s: p=0.01, n=31. f) Splitting the brain into six fragments inhibits foraging behavior one hour after incision, with worms unable to detect rotifers embedded in agarose. Tukey post-hoc tests for mean distance from food at the end of the assay, 3x vs other treatments p<=0.02, n>=9 per treatment. All error bands represent standard error.

To conclusively test the idea that more tiles allow better estimation of prey position, and to identify any other subtle effects of amputation, we sought to challenge the worms in a more dramatic, and ethologically-relevant way. We studied the hunting behavior of worms pursuing a single free-moving prey (i.e. a brine shrimp) - a challenging task in which nearly all hunts end in failure to consume the prey within the duration of the assay (we recorded 1/12 successful captures among intact worms in one experiment). We jointly tracked the positions of the worm and the shrimp (Fig. 4c, Video S4). Shrimp had lower velocity autocorrelations when in the presence of a worm (Fig. S7c), suggesting that they actively evade being hunted. Similarly, worms displayed lower velocity autocorrelations when the shrimp was present (Fig. S7d), showing that they actively hunt escaping prey. Worms with half their heads amputated displayed similar dynamics, but had slightly lower velocity autocorrelations compared to intact worms, specifically during foraging (Fig. S7d). These data suggest that amputation has a subtle impact on how worms hunt, but has little impact on the worms’ ability to detect stimuli or execute movements. We therefore asked whether amputated worms are worse at hunting.

We found that worms were able to track shrimp, moving towards the trajectory of the shrimp, with the worm lagging the shrimp by 20-30 seconds (Fig. 4c,d; Video S4). Worms missing half their heads were poorer at tracking shrimp (Fig. S7e). Next, in another cohort of worms, we removed increasing fractions of brain tissue and analysed postural dynamics and hunting behavior. This experiment, done in a richer behavioral setting, confirmed our previous finding that all postures and postural sequences are robust to large-scale amputation, and can be generated by arbitrary brain regions (Fig. S7f-j). Indeed, relative occupancy of postural states, and transitions between postures, were also largely indistinguishable between intact and amputated worms (Fig. S7i,j, Table S5,S6). However, we found that worms missing greater fractions of brain tissue displayed poorer tracking (Fig. 4d). In principle, this could be explained by the fact that some amputations also remove fractions of their mouth, and could reduce chemosensory sensitivity. To control for this possibility, we repeated this experiment, removing varying fractions of brain tissue but leaving the entire mouth intact in all cases. Again, worms missing greater fractions of brain tissue displayed poorer prey tracking (Fig. 4e), confirming that prey tracking is sensitive to the loss of brain tiles downstream of sensation. These data show that more brain tiles allow greater perceptual resolution.

Finally, we asked how brain tiles collectively generate coherent organismal behavior. One possibility is that brain tiles compute independently, with the motor systems effectively summing their separate outputs: i.e., the *H. miamia* brain would perform degenerate computations. Alternatively, it is possible that tiles interact, allowing computations to emerge from their collective activity: distributed computation. To test this hypothesis, we split heads into increasing numbers of fragments. Each fragment remained attached to the body; i.e. the number of brain tiles stays constant. We reasoned that this would disrupt interactions between brain tiles, but allow them to generate motor outputs independently. Consistent with this hypothesis, worms with their heads split into two or four fragments foraged similarly to intact worms, but worms with their heads split into six fragments did not (Fig. 4f). These worms were capable of coordinated movement, indicating that this defect was not mechanical (Video S5).

Together with our anatomical data showing extensive lateral connectivity (Fig. 1d; Fig. S1d), this suggests that circuit tiles do indeed interact, with coherent organismal behavior emerging from the collective activity of interacting brain tiles.

## Conclusion

Nearly a century ago, Lashley lesioned cortical regions from the brains of rats, and found that they could still perform spatial navigation and solve mazes^62,63^. From these and earlier experiments, he devised principles for brain organization: the principle of equipotentiality (effectively: brain regions are computationally pluripotent - or equipotent) and the principle of mass action (effectively: more brain is better). However, further work showed that Lashley’s principles do not accurately describe the organization of mammalian brains, which are indeed regionalized^13,64–66^. Strikingly, our results show that *H. miamia*’s diffuse brain, unlike typical bilaterian brains, does obey Lashley’s principles. We find that the *H. miamia* brain is not regionalized anatomically, cellularly, or functionally, and that all brain regions appear to perform most or all neural computations. We propose that the brain is composed of tiles of pluripotent neural circuits, and that these tiles interact to implement distributed computations. More generally, our results suggest that the space of realized brain forms may be broader than was previously assumed; brains do not have to be highly regionalized or stereotyped.

The precise historical sequence of the transition from nets to brains remains unknown, and is highly debated^9,10,67^. It has been proposed that ancient sensory organs connecting to diffuse nets served as nucleation centers for nascent brains^9,68^. In this view, the first brains would have been at least somewhat regionalized internally. Our work raises another possibility. We suggest that diffuse brains lack regionalization, and that the first brains may have retained the ancestral lack of regionalization and stereotypy that typifies diffuse nets. Perhaps these nascent brains then evolved elaborations for dense connectivity, new molecular signaling systems^59^, and possible new neural morphology and circuit motifs^46,69^ - coalescing in new behavioral affordances. Regionalized, stereotyped circuits may have evolved later, bringing forth the ground plan that enabled expansions into a diversity of bilaterian brains and a corresponding range of complex behavioral repertoires.

## Acknowledgments

We thank Ugne Klibaite, Brandon Logeman, Aravinthan Samuel, David Palmer, Ishaan Chandok, Aditi Chandra, Benjamin de Bivort, Madeleine Snyder, Julian Smith III, and the entire Srivastava Lab for helpful discussion. We thank D. Marcela Bolanos, Samantha Tseng, and Julian Kimura for assisting with data collection, Maria Ericsson and the Harvard Medical School Electron Microscopy Core Facility for high-pressure freezing and sample preparation, Ed Soucy and Jazz Weisman for behavioral advice, and Tim Sackton and Tommy Tang for single-cell analysis advice. VC and APK are Fellows of the Jane Coffin Childs Fund for Medical Research. VC acknowledges funding from the Harvard Brain Initiative’s Postdoctoral Pioneer Award. AS was supported by a grant from the Harvard Museum of Comparative Zoology. This project was supported by an NIH BRAIN R34 award (1R34DA061984) to MS.

## Author Contributions

VC and MS designed the project. VC performed behavioral experiments. VC and MRN performed computational behavioral analyses. VC, APK, AS, KAPH, and RS performed molecular labeling and imaging experiments. VC analyzed transcriptomic and imaging data. JL enabled electron microscopy. MS and LM supervised the project. VC and MS wrote the manuscript with input from all authors.

## Methods

### Animal husbandry

Juvenile *Hofstenia miamia* worms were reared in communal plastic tanks with artificial sea water (37ppt, pH between 7.8 and 8.2), held at room temperature with an approximate day-light cycle. Sea water was replaced twice weekly, and the worms were fed marine rotifers (*Brachionus plicatilis*).

For behavioral experiments, worms were isolated from their culturing tanks, and placed in individual wells of 6- or 12-well plates (Falcon #353046). Worms were typically 3-5 weeks post hatching, at which point they varied in their sexual maturity. Preliminary observations indicated that this variation did not affect foraging behavior, and each experiment was initiated with worms from the same communal tank, of approximately the same age and size. Worms that were in the middle stages of sexual maturity were fed rotifers, brine shrimp (*Artemia sp.*) or a mix of the two, as previously described^42^. To amputate worms, we anesthetized them in 15% tricaine (ethyl 3-aminobenzoate methanesulfonic acid; Sigma #E10521) for 10-20 minutes, and amputated them with micro knives (Fine Science Tools #10316-14) before washing in artificial sea water to allow recovery. Control worms in these experiments were anesthetized for the same amount of time, but were not amputated. For foraging experiments, worms were starved for 1-2 weeks.

### Dyes and live imaging

We used several live dyes to stain *H. miamia* brains. We used the voltage dye DiBAC4(3) (Invitrogen #B438) to stain the nervous system. We dissolved DiBAC4(3) in pure ethanol to create a 1mg/ml stock solution. We diluted this 1:1000 in artificial sea water with 15% tricaine. Worms were incubated in this solution for 30 min. They were then mounted in a gel of 2% methylcellulose (in artificial sea water containing 15% tricaine) on a glass slide, compressed under a glass coverslip with clay feet, and imaged. The calcium dye ICR-1 AM (Ion Biosciences #1091F) was reconstituted in DMSO and then diluted in artificial sea water with 15% tricaine to a concentration of 40µM. Worms were incubated in this solution for 1-2hrs, then mounted on a glass slide in methylcellulose, using the same procedure as above. SiR-Tubulin (Cytoskeleton, CY-SC002) was dissolved in 50µL DMSO according to manufacturer instructions, then worms were incubated in a 1:1000 solution in artificial sea water with 15% tricaine for 90m prior to mounting as described above. *Troponin::Kaede* animals, described previously^51^, were anesthetized in tricaine, and mounted using the same procedures as above.

### Irradiation

To kill adult pluripotent stem cells, seven days before amputation, worms were exposed to gamma radiation for a total dose of 10000 rad. Worms were then amputated and their behavior was assayed two days later as described below. All worms survived irradiation.

### Immunofluorescence

We performed immunofluorescence using previously described protocols^42^. Briefly, worms were fixed in 4% paraformaldehyde in artificial sea water for 1h at room temperature. Fixed animals were washed in PBST (PBS + 0.1% Triton-X-100) three times, then blocked in 10% goat serum (ThermoFisher, #16210072) in PBST for 1h at room temperature. Samples were incubated in primary antibodies in blocking solution for 48h at 4°C, washed thoroughly (6x 20 minutes) in PBST, incubated in blocking solution for 1h at room temperature, then incubated in secondary antibody overnight at 4°C. Animals were washed thoroughly once more (6x 20 minutes) in PBST, then counterstained with Hoechst 33342 and SiR-actin (where applicable). All steps were carried out on an orbital shaker. Primary antibodies used: Par3 (St. John’s Laboratory #STJ94951, 1:200), β-catenin (Fisher Scientific #RB9035P0, 1:200), pERK (Cell Signaling Technologies #4370T, 1:200) ^70^, FMRFamide (EMDMillipore #AB15348, 1:1000)^36^.

Counterstains used: Hoechst 33342 (ThermoFisher #62249, 1:800) and SiR-actin (Cytoskeleton #CY-SC001, 1:1000)^50^. Goat anti-Rabbit IgG (H+L) Cross-Adsorbed Secondary Antibody, Alexa Fluor™ 568 (ThermoFisher #A-11011, 1:800) was used as a secondary antibody.

### Electron Microscopy

We fixed hatchling juvenile worms using high-pressure fixation. A mix of intact worms and amputated heads in 15% tricaine in artificial sea water were placed in 100µm-tall aluminium planchettes, and were rapidly frozen in liquid nitrogen using a Leica EM ICE high-pressure freezing instrument. These planchettes were then transferred under liquid nitrogen into cryovials containing a freeze substitution cocktail (1% OsO4 and 0.1% uranyl acetate). Freeze substitution involved holding the cryovials at -90°C for 36-48h, followed by warming up to -60°C over 6h and holding for 6h, warming up to -30°C over 6h and holding for 3h, warming up to 0°C over 6h and holding for 1h, and finally warming up to room temperature over 4h. Specimens were then rinsed in acetone three times for 10m each. They were then extracted from the planchettes and embedded in an Epon-Araldite resin. Resin blocks were baked at 60°C for 48h, after which they were sectioned with a Leica UC6 ultramicrotome using a DiATOME diamond knife, and collected onto tape using ATUM^71^. Sections of tape were then mounted on silicon wafers as previously described^71^, and were imaged using a Zeiss Sigma scanning electron microscope. Neurites in Fig. 1j,k were segmented using Cellpose-SAM^72^ with manual correction.

### Riboprobe synthesis and fluorescence *in situ* hybridization (FISH)

We performed FISH using protocols described previously^31,73^, with minor modifications. Briefly, we used nested PCR to extract template sequences from cDNA. Riboprobes were synthesized from these templates using a digoxigenin (DIG) labeling mix, as previously described^31,73^.

Worms were starved for 1-2 weeks prior to fixation to ensure that their guts were empty. They were then fixed in 4% paraformaldehyde in artificial sea water for one hour at room temperature. We washed worms in PBST (PBS + 0.1% Triton-X-100), 50% PBST/50% MeOH solution, and chilled MeOH for 5m each. They were then transferred to baskets in 24-well plates containing 800µL of solution per well, washed once in 50% MeOH/50% PBST, then twice in PBST. To remove pigmentation and autofluorescence, they were bleached in a solution of 88.5% MilliQ water, 4% hydrogen peroxide, 2.5% 20X SSC, and 5% formamide, then washed in PBST twice. Worms were then incubated for 10 minutes in Proteinase K solution (0.1% SDS, 0.9% MilliQ water, and 0.01% Proteinase K in PBST), post-fixed in formaldehyde solution (4% formaldehyde in PBST) for 10m, and washed in PBST. They were then incubated in 50% PBST/50% PreHyb (50% DI Formamide, 25% 20X SSC, 0.5% Tween-20, 24.5% MilliQ water, 0.001g/mL Yeast RNA), transferred to 100% PreHyb and incubated at 56°C for 2hrs. 1µL of probe was then mixed with 799µL of hybridization solution (50% DI Formamide, 25% 20X SSC, 0.5% Tween-20, 5% dextran sulfate, 19.5% MilliQ water, 0.001g/mL Yeast RNA), denatured at 72°C for 5m, and distributed in the wells. The worms were moved into the probe solution, sealed, and incubated at 56°C overnight. The following day, the worms were washed twice in PreHyb, twice in 50% PreHyb/50% 2X SSCT (10% 20X SSC, 0.1% TritonX-100, 89.9% MilliQ water), 100% 2X SSCT, then 0.2X SSCT (1% 20X SSC, 0.1% TritonX-100, 98.9% MilliQ water), for 30m each at 56°C. They were then cooled at room temperature for 10mins, washed in PBST, incubated in blocking solution for 1hr (90% PBST, 5% horse serum, and 5% 10X Casein), then moved to 1:1500 blocking solution:Anti-DIG, sealed, and left at 4°C overnight. The next day they were washed six times for 20m each in PBST, incubated in tyramide buffer for 10m, then development solution (0.1% rhodamine, 0.1% IPBA, and 0.01% H2O2 in tyramide buffer) for 10m. They were then washed in PBST 3 times before being moved to a solution of 1:500 Hoechst 33342 (ThermoFisher #62249) in PBST, sealed, and left at 4°C overnight. Finally, they were washed in PBST, and then mounted on glass slides in VECTASHIELD® PLUS Antifade Mounting Medium (Vector Laboratories, H-1900). All steps except the 56°C incubations were carried out using an orbital shaker.

### FISH analysis

We collected all images with a Leica SP8 point-scanning confocal microscope. 3D images of FISH-labeled neural markers in brains were generally acquired through a 20x objective. We then generated a maximum intensity z-projection of the brain for each of these images, and trained a supervised pixel classifier in ilastik to segment pixels that had FISH signal. These binary images were then smoothed in FIJI through a process of erosion and hole filling, before computational analysis.

To quantify regionalization of neural markers within the brain, we computed two indices of regionalization. First, we partitioned each brain into a heatmap containing 25 pixels, and we calculated the proportion of pixels containing fluorescent puncta. We then calculated the Gini coefficient (a standard measure of inequality) of each heatmap. To correct for differences in regionalization expected as an artifact of sparseness, we simulated brains with uniformly distributed fluorescent puncta of the same sparseness. Computing the difference in Gini coefficients between real and matched simulated brains provided a robust index of regionalization. Second, for each brain we constructed the convex hull of the set of fluorescent puncta positions, treating puncta as points in the imaging plane. We computed the area enclosed by this convex hull and normalized it by the area of the polygon defining the brain boundary, yielding a dimensionless coverage ratio that quantifies the spatial extent of puncta relative to the entire brain. To generate a second index of regionalization, we measured the difference between real ratios and corresponding ratios within simulated brains with uniformly distributed puncta.

### Behavior

#### Behavioral imaging

All behavioral experiments were conducted in 6-well plates filmed with 850nm infrared side-lighting from two LED strips behind a white acrylic diffuser (McMaster-Carr #8505K741), within a custom behavioral chamber shielded from visible light. The chamber was cooled with a USB fan to maintain temperatures between 22 and 25°C. Experiments were filmed in parallel using four machine vision cameras (FLIR Blackfly BFS-U3-120S4M-CS and Basler Ace ac4024-29um USB 3.0 monochrome, with Tamron M118FM16 16mm lenses) at 12 megapixels and 10Hz for 1h, streaming to a custom-built desktop computer; each video contained four or six wells of a single plate within its field of view. We then used a custom MATLAB script for pre-processing: background subtraction, contrast enhancement, and cropping the video into a set of separate output videos for each well of the plate. We typically scaled videos to a resolution of 1000x1000 pixels, masking areas outside the arenas, and aligning their centers to make xy coordinates comparable across videos.

#### Immobilized rotifer foraging assay

To study the foraging behavior of *H. miamia* presented with immobilized prey, we developed an assay to present worms a choice between blocks of agarose that either did or did not contain groups of live, immobilized rotifers. Rotifers were embedded in ∼1% agarose blocks using the following protocol. For each agarose block, 100µL of a well-mixed stock solution of rotifers (on the order of thousands of rotifers per mL) or 100µL of artificial sea water for controls, was first aliquoted into the bottom of a subset of wells of a 96-well plate. 100µL of hot 2% UltraPure Low Melting Point agarose (Invitrogen #16520-100) was then added immediately above. The plate was then cooled at 4°C for 30m. Each solid agarose block was then placed against the wall of a well of a 6-well plate, with a block of agarose alone and a block of agarose containing rotifers at opposite ends of the wall. A small volume of 2% agarose was then pipetted onto this block to glue it to the wall of the plate. 5-10m later, 6ml of artificial sea water was pipetted into the well, a single *H. miamia* worm was introduced to the well and the behavioral experiment initiated.

#### Tracking and foraging analysis

We trained custom models in DeepLabCut^74^ and SLEAP^75^ to track keypoints on the worm. To study foraging behavior, we quantified the position and movement of worm centroids inferred from these keypoints. We systematically excluded worms from the analysis if they were tracked poorly, or which were inactive for sustained periods over the course of the experiment (a minority of worms fit these exclusion criteria across our experiments, independent of their feeding or amputation status). To quantify the proportion of time spent near food (i.e. rotifers embedded in agarose), we defined an experiment-specific threshold distance, and quantified the proportion of frames in which the worm was within that distance.

#### Postural analysis

For the datasets used for postural analysis, we tracked seven keypoints along the midline of the worm. We smoothed the xy coordinates of these points, used median-filling to remove NaNs, and then fit a 20-jointed spline to this data. Preliminary observations of the worms suggested that their postural dynamics are well represented by three features: the curvature of their midlines, their body lengths, and the speeds the worms travel at. We calculated the curvature of the spline as the normalized ratio of the sum of the lengths of its segments to the length between the first and last points. This ratio was scaled to lie within the range [-1,1], where -1 represented a sharp left bend, 0 represented the worm’s posture when straight, and 1 represented a sharp right bend. Worm length was calculated as the sum of the segments composing the spline; this was z-scored to enable comparison across worms and across experiments in which the measured length varied slightly as a function of precise camera parameters. Speed was calculated as minimum displacement between splines in consecutive frames. All measures were smoothed and filtered to remove outliers. We used PCA to reduce this curvature-length-speed space to 2D; we then used k-means clustering to partition this postural space into elemental postures as described in the text. We used multiple large language models to assist with generating analysis code.

#### Single-cell RNA sequencing analysis

We integrated single-cell RNA sequencing libraries from three previous sequencing experiments. These included worms that ranged from hatchlings (i.e. juvenile worms that have just emerged from their egg shells) to adults, and worms in various stages of regeneration after a transverse amputation. We re-aligned these libraries to an updated transcriptome^58^. We then regressed library identity to correct for batch effects, identified neural clusters, extracted and re-clustered neural cells.

#### Hunting behavior assay and analysis

To study the behavior of *H. miamia* actively hunting escaping prey, we introduced a single worm and a single brine shrimp to each well of a 6-well plate. We filmed these arenas for 1h at 10Hz in infrared lighting conditions, using the imaging parameters described above.

We then tracked worms and brine shrimp using custom-trained SLEAP models. To find the velocity of the worm at frame i while smoothing tracking noise, we averaged the center-of-mass velocity over five frames before and after frame i. The velocity correlation function was then calculated for the entire trajectory of each sample using the definition 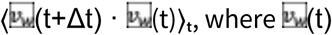 denotes the velocity of the worm at time t, and the average is taken over time t. The final velocity correlation function shown in Fig. S7d is obtained by averaging the velocity correlation functions of worms across different experiments with the same treatment.

To quantify the correlation between the velocity of the worm and the trajectory of the shrimp (Fig.4d,e; Fig. S7e), we used the smoothed worm velocity vᵥ and calculated its correlation with the vector connecting the worm’s position to the shrimp’s position in earlier frames. The correlation is defined as 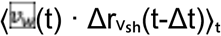, where Δrᵥₛₕ(t-Δt) is the unit vector pointing from the worm at time t to the shrimp at time t-Δt.

