## Supplementary Materials for "Distributed neural computation and the evolution of the first brains"

| Cluster | Speed | Curvature | Length |
|---|---|---|---|
| 0 | 0.42 | -0.00 | -0.06 |
| 1 | 0.03 | -0.73 | 1.19 |
| 2 | 0.04 | 0.78 | 0.06 |
| 3 | 0.04 | -0.01 | -0.70 |
| 4 | 0.04 | -0.00 | -0.12 |
| 5 | 0.02 | 0.68 | 1.84 |
| 6 | 1.02 | -0.00 | -0.01 |
| 7 | 0.04 | -0.79 | -0.06 |
| 8 | 0.02 | -0.61 | 2.66 |
| 9 | 0.07 | -0.01 | 0.32 |

**Table S1:** Median values of normalized speed, normalized curvature, and normalized length for postural clusters from Fig. 2h.

Tables S2-S6, and Videos S1-S5 provided as supplementary files.

**Table S2:** Statistical differences in postural cluster occupancy between amputated and intact worms foraging for immobilized rotifers.

**Table S3:** Statistical differences in postural cluster transition frequencies between amputated and intact worms foraging for immobilized rotifers.

**Table S4:** List of genes and primer sequences for nested PCR used for fluorescence *in situ* hybridization.

**Table S5:** Statistical differences in postural cluster occupancy between amputated and intact worms hunting actively-escaped brine shrimp.

**Table S6:** Statistical differences in postural cluster transition frequencies between amputated and intact worms hunting actively-escaped brine shrimp.

**Video S1:** Confocal z-stack of superficial tissue in the head of a worm stained with a calcium dye (ICR-1 AM). Dense peripheral neural processes are visible. Scale bar: 50μm

**Video S2:** Videos of representative fed and starved worms foraging, when presented a choice between rotifers immobilized in agarose (right side of dish) and plain agarose (left side). DeepLabCut tracking is shown below the raw videos. All videos sped up ~100x.

**Video S3:** Worm missing the majority of its head, one hour after amputation, can glide, can generate left and right turns, and does not display obvious behavioral defects. Video in real time.

**Video S4:** Joint tracking of worm and a brine shrimp during a hunt. Video sped up 20x.

**Video S5:** Worm with head split into six fragments is also capable of coordinated movement. Video in real time.

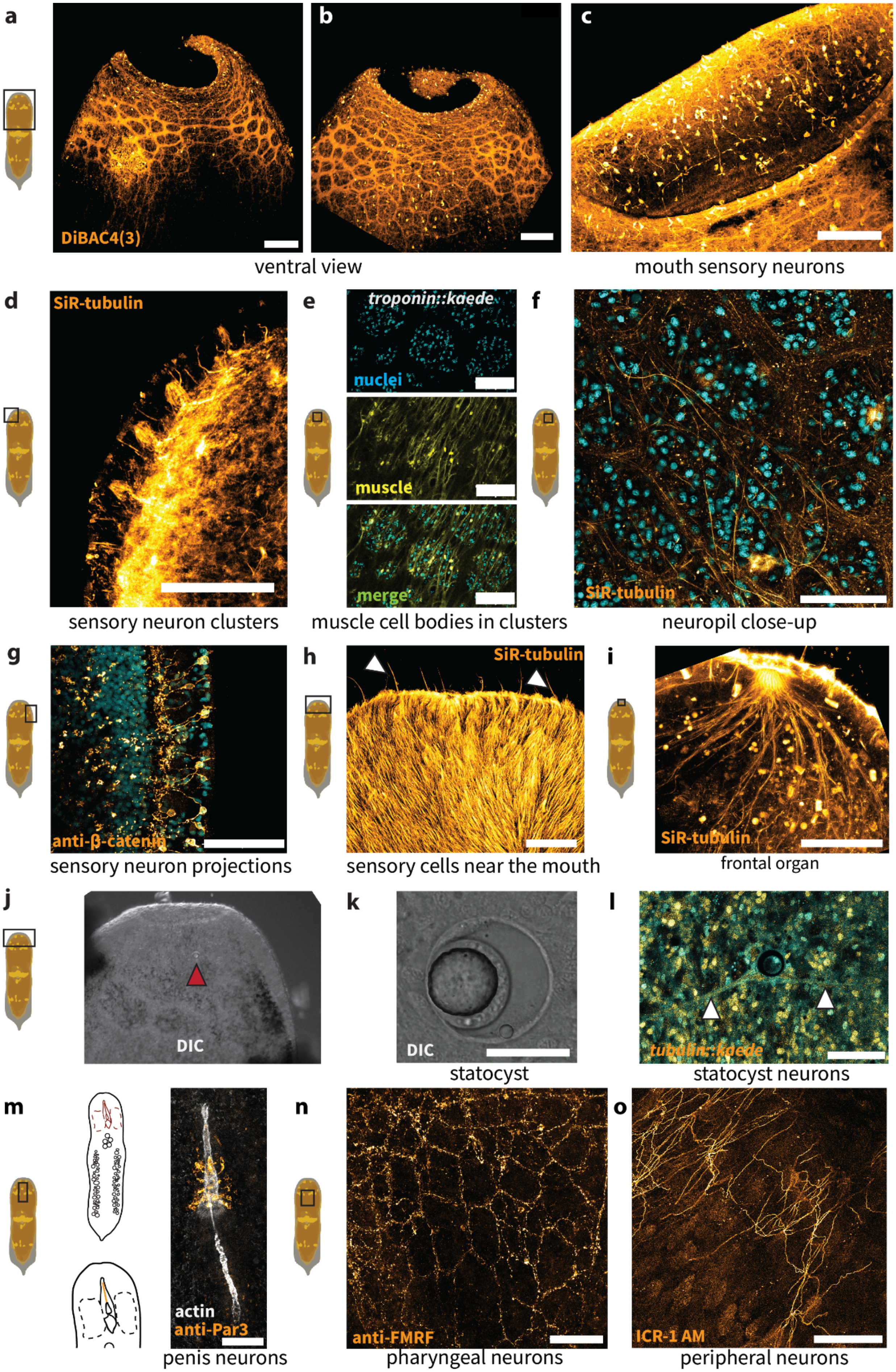

a
DiBAC4(3)
b
c
ventral view
mouth sensory neurons
d
SiR-tubulin
e
troponin::kaede
nuclei
muscle
merge
f
SiR-tubulin
sensory neuron clusters
muscle cell bodies in clusters
neuropil close-up
g
anti-β-catenin
h
SiR-tubulin
i
SiR-tubulin
sensory neuron projections
sensory cells near the mouth
frontal organ
j
DIC
k
DIC
l
tubulin::kaede
statocyst
statocyst neurons
m
actin
anti-Par3
n
anti-FMRF
o
ICR-1 AM
penis neurons
pharyngeal neurons
peripheral neurons

**Figure S1: The structure of the *H. miamia* nervous system.** a, b) Ventral view of the brain, stained with the voltage dye DiBAC4(3), reveals that the neuropil typically wraps around the head, with a lower density of 'edges' of neuropil at the ventral midline. These images also show that the specific shape of neuropil varies across animals. c) Close-up of mouth, stained with DiBAC4(3), reveals putative sensory neurons arranged around the mouth. d) Tubulin stain reveals clusters of superficial sensory neurons (likely H1). e) Fluorescent peripheral muscle cells (seen in yellow) in a transgenic *troponin::kaede* worm have cell bodies that lie within the cellular clusters (nuclei stained with Hoechst and labeled in cyan) embedded in the neuropil. f) Staining with a tubulin dye reveals long neural projections within neuropil. g) Immunostaining against β-catenin reveals sensory neurons embedded in the skin and projecting into the neuropil. h) Exterior of worm labeled with tubulin dye reveals dense epidermal cilia, and sensory cells with long cilia near the mouth, visible on the anterior (white arrows). i) The frontal organ, putatively a sensory and secretory structure dorsal to the mouth[36], is labeled with a tubulin dye, showing long projections extending into the brain. j) View of worm head under DIC, with the statocyst labeled (red arrow). k) Close-up view of the statocyst, putatively a gravity-sensing organ, viewed under DIC, shows a cell within the organ that contains a mineralized statocyst, neighbored by two peripheral parietal cells. l) Labeling neurons in a photoconvertible (*tubulin::kaede*) transgenic line reveals two (or possibly a few) neurons (white arrows) that appear to innervate the statocyst, forming a structure previously referred to as the 'dorsal commissure'[36]. Image courtesy of Julian Kimura. m) Immunostaining against Par3 reveals a population of neurons that innervate the penis. The schematic on the left shows a ventral view of an adult worm, with an enlarged view of the male copulatory apparatus depicted below (penis highlighted)[42]. n) Staining with an anti-FMRFamide antibody reveals a sparse network of neurons surrounding the pharynx, similar to the posterior nerve net and consistent with previous descriptions. o) Staining with a calcium dye reveals dense peripheral neural projections, possibly innervating the skin. Scale bars: 100μm (a,b,c), 50μm (d,e,f,h,i,l,n,o), 20μm (g,k).

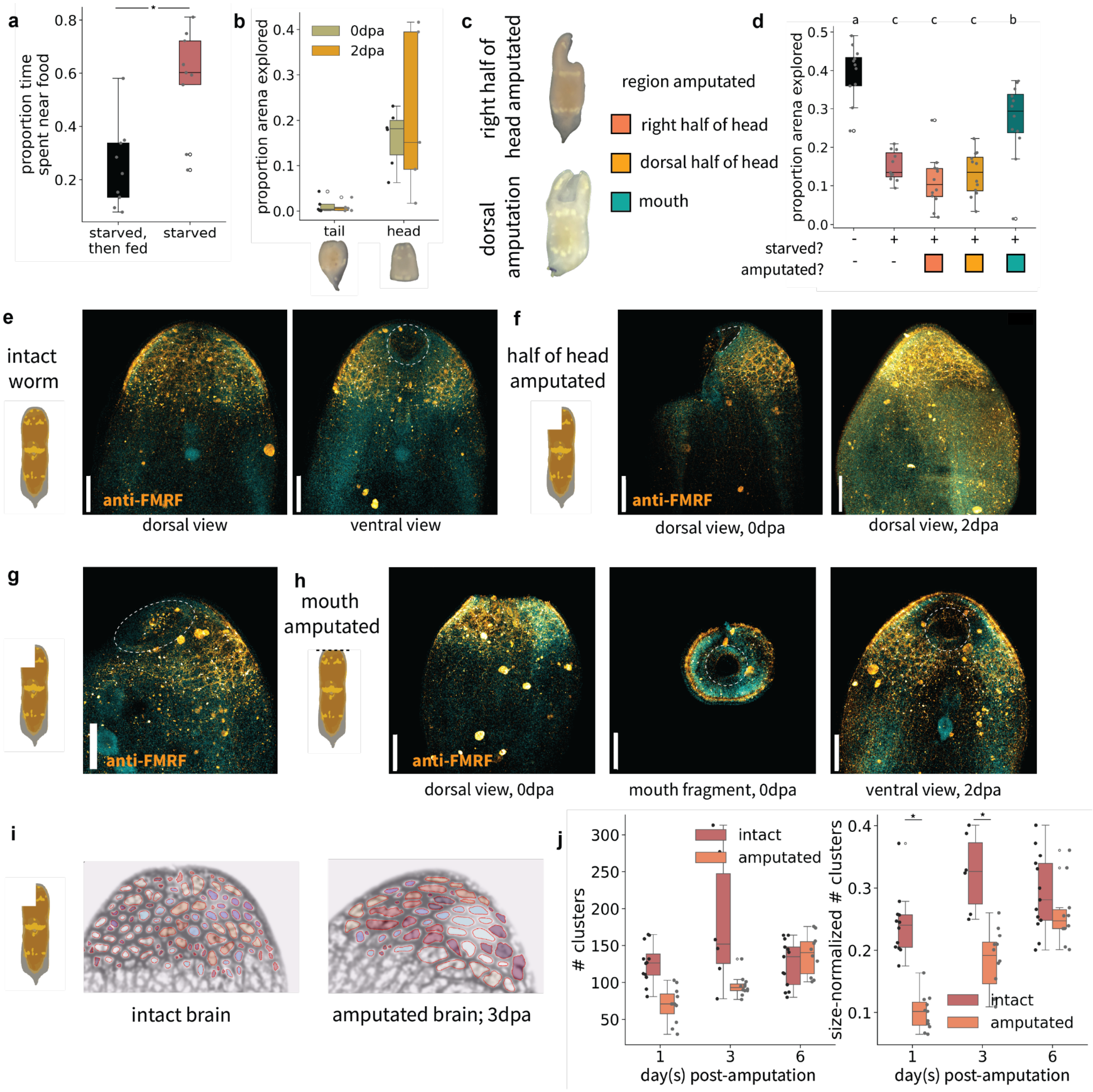


**Figure S2. Visualizing the anatomical and behavioral consequences of amputations of brain and sensory regions in *H. miamia*.** a) Previously starved worms, when fed for 6h, cease foraging, showing that hunger in *H. miamia* is assessed and updated frequently (likely continuously). t-test p=0.001, n=9 per treatment. b) Tail fragments cannot move in a coordinated manner, but head fragments do - and forage normally (see Fig. 2d). c) Representative images of worms after amputation of the right and dorsal half of the head; background removed and contrast adjusted. d) These worms do not have movement defects, and do not explore the arena differently from intact controls. However, fed intact worms, and starved worms missing their mouths, explore substantially more of the arena, consistent with their displayed lack of foraging (Fig. 2e); Tukey post-hoc p<0.002 for relevant comparisons. Staining with an antibody against FMRFamide shows the effects of amputation. Intact worms visible in (e), worms missing half their heads in (f, g), and worms with mouths amputated in (h). g) Two days after amputation of half the head, a circular mouth is visible. This indicates that the wound closes through a process in which the dorsal and ventral wound edges meet. i) Segmentation of cell clusters within neuropil allows quantification of the effects of amputation; here, a representative image of an intact worm (left image), and a worm missing half its head (right image) 3 days after amputation are shown. Amputated worms have fewer cell clusters. j) The number of clusters in a worm missing half its head is reduced compared to intact controls 1 ($p<0.0001$, $n>=10$) and 3 ($p<0.0001$, $n>=6$) days after amputation, but regenerates by 6 ($p=0.18$, $n>=10$) days. Scale bars: 200μm (e-h)

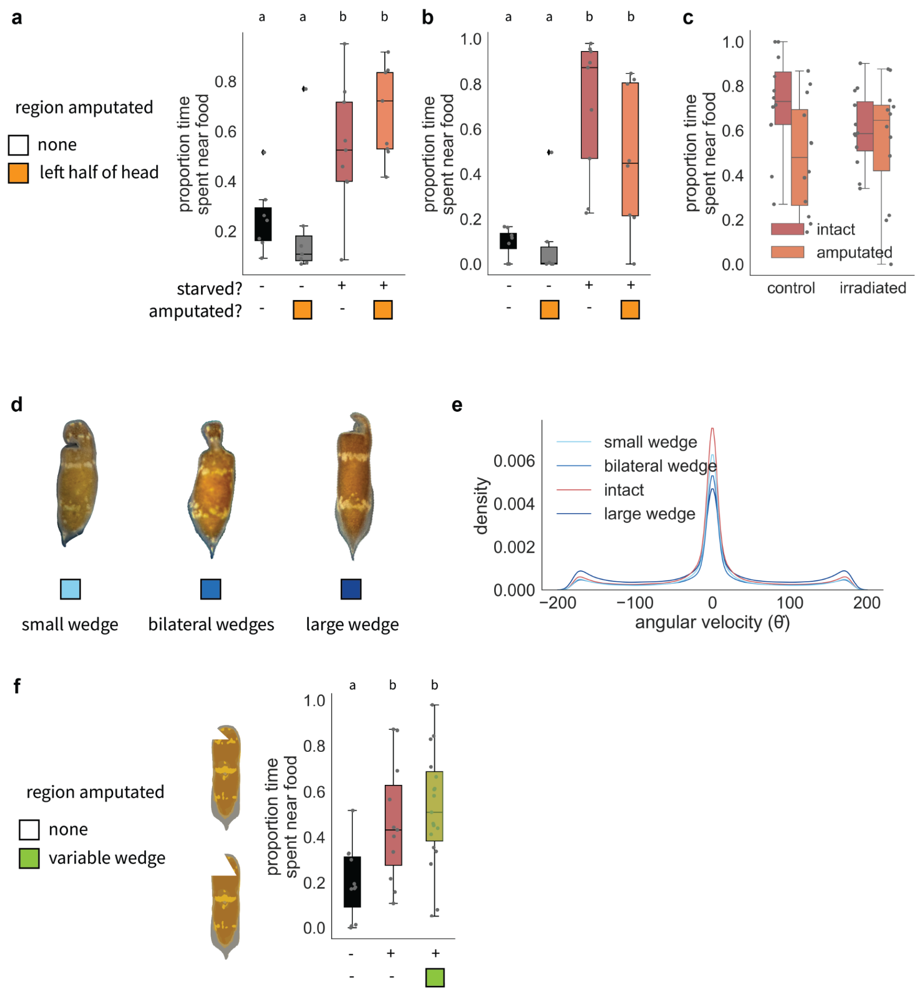


**Figure S3. Behavioral robustness to amputation of brain regions.** Amputation of the left half of the head does not affect foraging behavior, either two days after amputation (a), or immediately after (b). Pairwise t-tests, FDR-corrected $p<0.05$, $n>=6$ per treatment in each experiment. c) Irradiated worms (lacking stem cells that allow regeneration) continue to forage normally, both when their brains are intact and when the right half of their head has been amputated. Tukey post-hoc tests $p>0.05$, $n=12$ per treatment. d) Representative images of wedge amputations in Fig. 2g; background removed and contrast adjusted. e) Kernel density estimate of angular velocity distributions, showing no major differences as a result of amputation, and no apparent turn bias. Amputation of wedges of varying size, ranging from ~10% to ~80% of the brain, has no detectable effect on foraging behavior (f). Pairwise t-tests, FDR-corrected $p<0.05$, $n>=11$. Letters above boxplots indicate significance levels after pairwise tests with multiple testing correction.

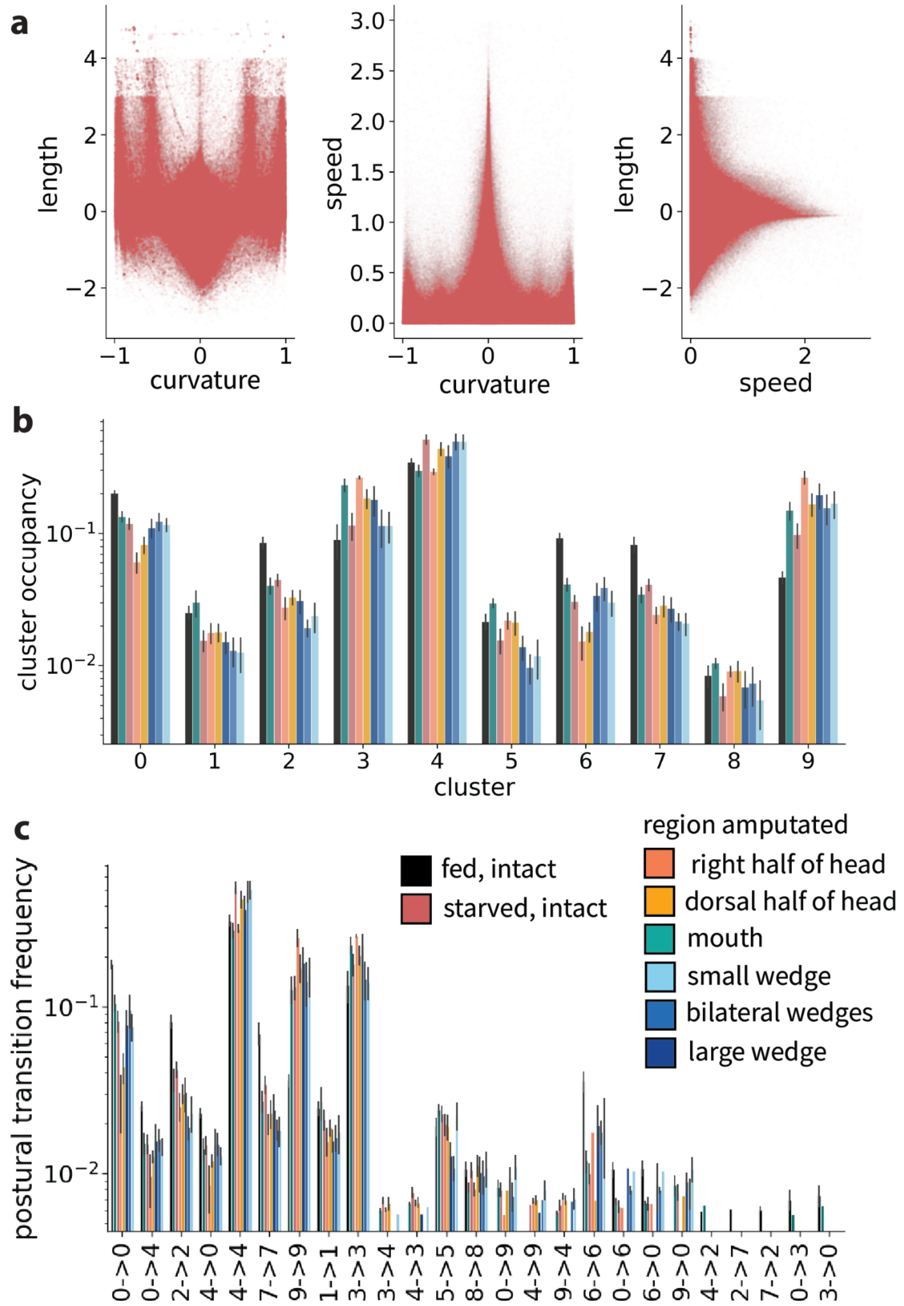


**Figure S4: Postural dynamics are robust to arbitrary amputations of brain tissue.** a) Postural space for all worm data, showing relationships between length, curvature, and speed. All measures are normalized and filtered; see Methods for details. b) Worms occupy all postural clusters regardless of amputation treatment. c) Worms display an invariant set of transitions between postural clusters (i.e. behavioral sequences) regardless of amputation treatment. Treatment colors in (b) and (c) are the same as those in Fig. 2. See Tables S2 and S3 for statistics.

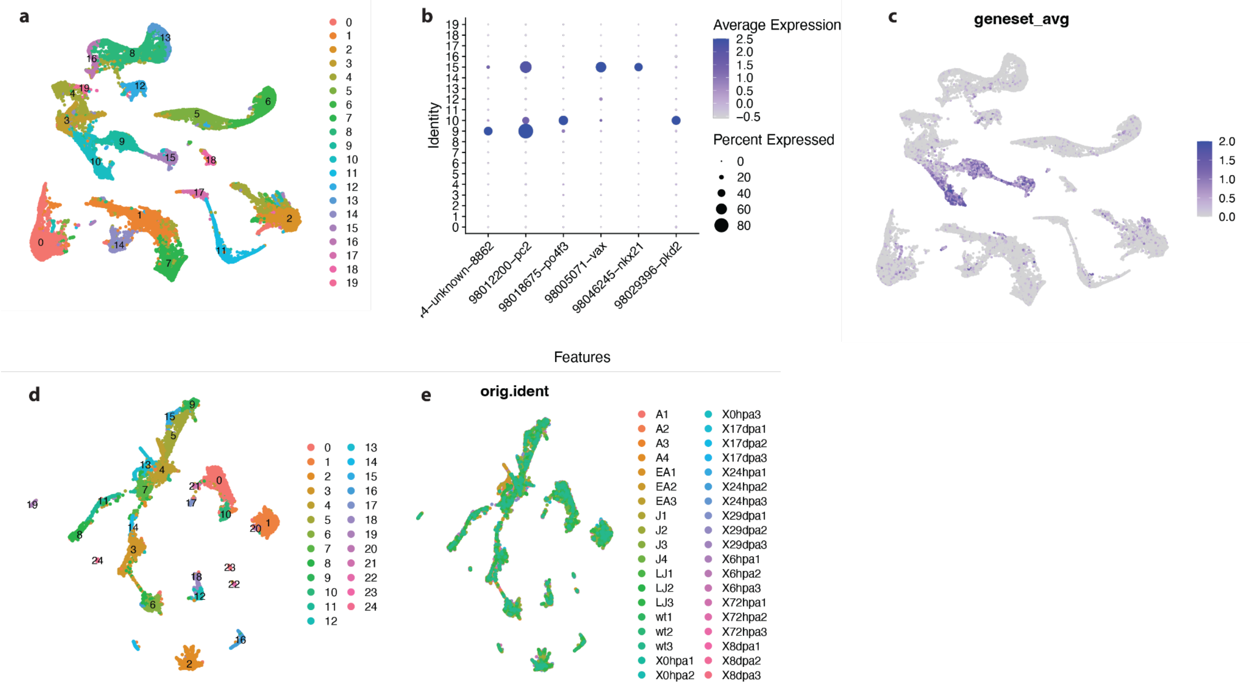


**Figure S5: Single-cell RNA sequencing analysis identifies neuron type markers.** a) UMAP of integrated single-cell data from[54,58], showing 20 clusters. b) Dotplot of previously-identified neural markers shows that they are specifically expressed in clusters 9, 10, and 15, suggesting that these clusters contain neurons. c) Mean expression of these six neural markers projected onto the UMAP shows that they specifically label clusters 9, 10, and 15. Subsetting and reclustering neurons shows 25 neural clusters (d), and that after integration and batch correction, each input library is represented in each cluster (e).

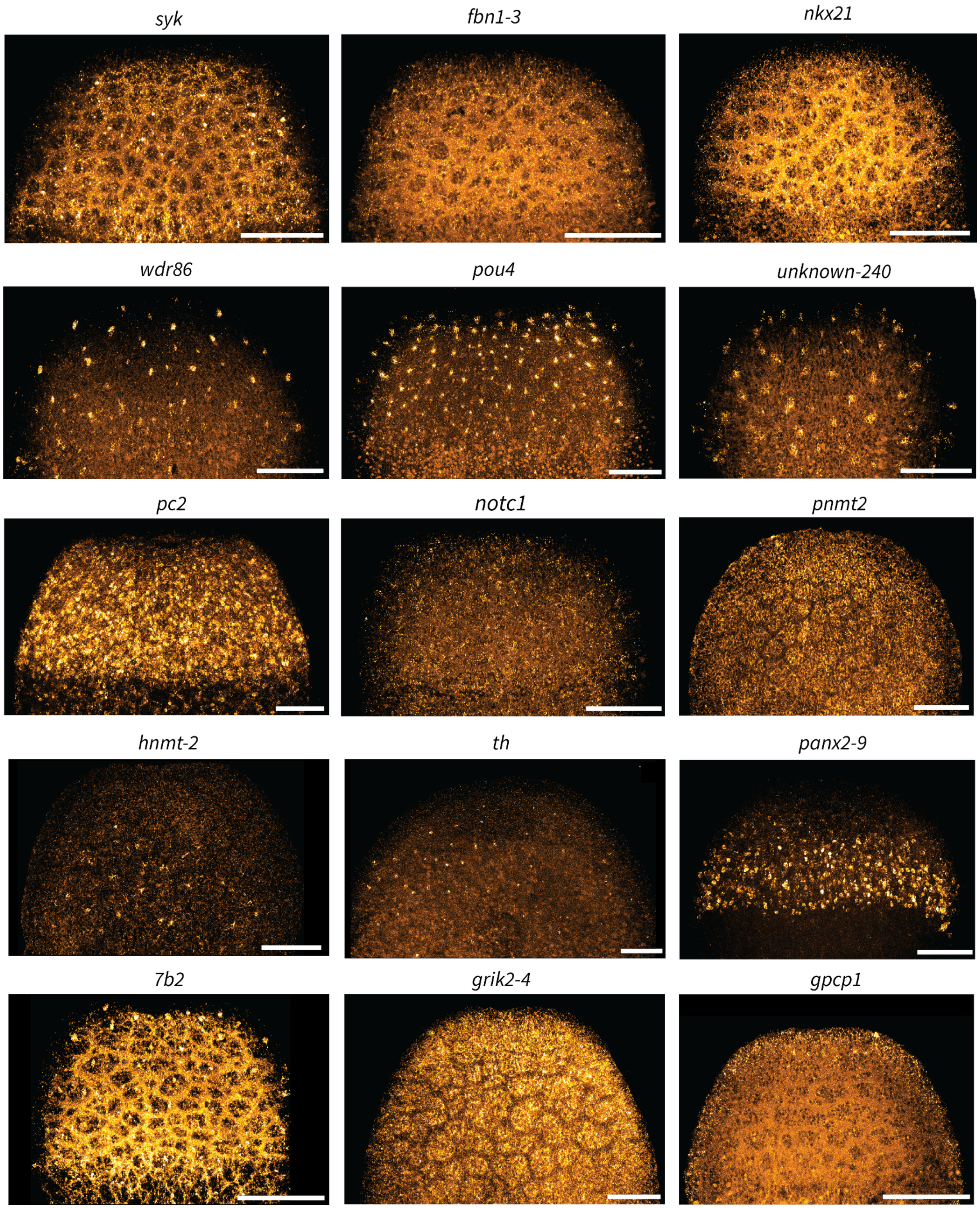


**Figure S6: Fluorescence *in situ* hybridization of neural markers.** Representative images of fluorescence *in situ* hybridization for neural markers, including re-imaging of previously-published staining (*syk, fbn1-3, nkx21, wdr86, pof43, unknown-240, pc2, notc1, th*)[54]. Scale bars: 100μm.

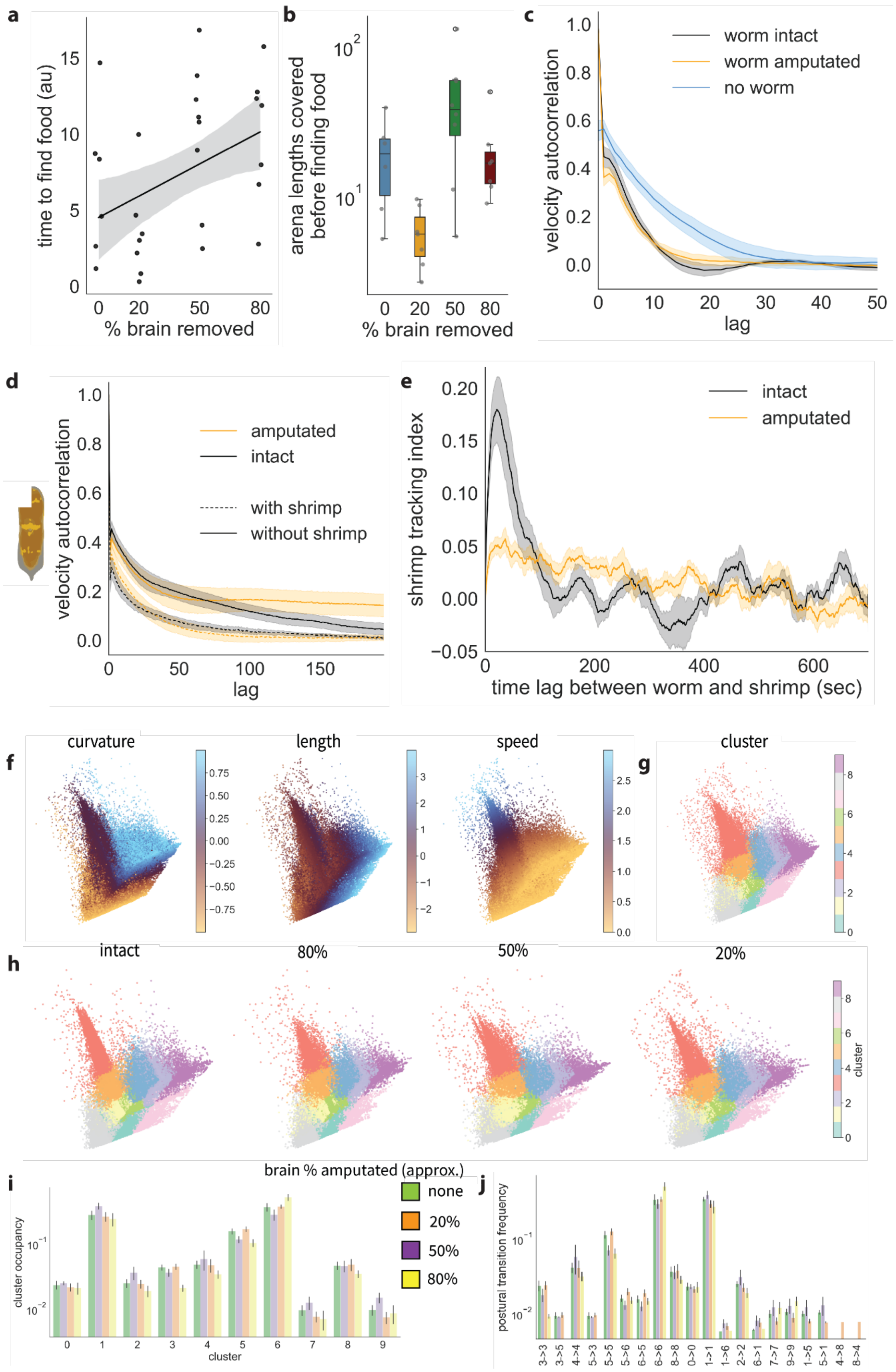
a
time to find food (au)
% brain removed
0
20
50
80
5
10
15
b
arena lengths covered before finding food
% brain removed
$10^2$
$10^1$
c
velocity autocorrelation
lag
worm intact
worm amputated
no worm
d
velocity autocorrelation
lag
amputated
intact
with shrimp
without shrimp
e
shrimp tracking index
time lag between worm and shrimp (sec)
intact
amputated
f
curvature
length
speed
g
cluster
h
intact
80%
50%
20%
cluster
i
brain % amputated (approx.)
none
20%
50%
80%
cluster occupancy
cluster
j
postural transition frequency
3->3
3->5
4->4
5->3
5->5
5->6
6->5
6->6
8->8
0->0
1->1
1->6
2->2
6->1
7->7
9->9
1->5
5->1
4->8
8->4

**Figure S7: Kinematic and postural analysis of worms hunting free-moving shrimp.** a) worms missing larger fractions of their brains take longer to find rotifers embedded in agarose. Linear regression p=0.02, n=28. b) The distance traveled by worms prior to locating the rotifers (quantified as approximate arena lengths covered) is not primarily a function of how much brain tissue they are missing. c) Shrimp have greater velocity autocorrelations in the absence of worms, suggesting that their behavior changes when they are being hunted. d) Worms have greater velocity autocorrelations in the absence of shrimp, suggesting more persistent movement. Moreover, amputated worms are less persistent in their direction of motion when hunting shrimp. e) Shrimp tracking index for intact worms, and worms missing half their heads shows that amputated worms are poorer at tracking shrimp. f) A 2D postural space constructed from splines fitted to worm midlines. g) k-means clustering of intact worm data within this space defines elemental postures. h) Mapping cluster labels onto amputated worms shows that, as in Fig. 2, amputation of brain regions does not affect the generation of elemental postures. i) Cluster occupancy is generally similar across amputation treatments. j) Transitions between postural states also display similar frequencies across amputation treatments. All error bands represent standard error.
